# Transcriptomic profiles from stereo-EEGs reveal the local cell microenvironment in human epilepsy

**DOI:** 10.1101/2025.09.16.676570

**Authors:** Julian Larkin, Anuj Kumar Dwivedi, Arun Mahesh, Albert Sanfeliu, Vijay Tiwari, Peter Widdess-Walsh, David C. Henshall

## Abstract

Objectives: Our understanding of the pathomechanisms of epilepsy has improved through techniques that access the living human brain. We recently reported that explanted stereo-electroencephalography (SEEG) electrodes from patients with epilepsy carry residual biomolecules and cells which may be utilised for transcriptome and DNA methylation profiling. Methods: Here, we applied bioinformatic and other analyses to explore the transcriptomes (RNA sequencing-based) of those SEEG cases to better understand the types of recovered transcripts in terms of representation of genes expressed by different cell types, brain structures, and the extent to which the signal may reflect local epileptiform activity. Results: Electrodes from all clinical cases retained protein-coding transcripts which reflected the local molecular microenvironment as well as epileptiform activity. Expression of genes involved in housekeeping functions as well as markers of neuronal activity were consistent between patients and between the electrode locations within the brain. We detected transcripts representing various cell types and subtypes including excitatory and inhibitory neurons, all major classes of glia, and endothelial cells, as well as transcripts enriched in specific brain regions. Several genes showed a gradient of expression depending on the electrode position within the brain. We found examples of gene expression that correlated with epileptiform activity as recorded by SEEG. Interpretation: These findings extend the evidence that SEEG electrodes reflect the molecular microenvironments of brain activity in patients with epilepsy, both at sites of seizure onset and within the wider seizure network. The approach has potential applications in intraoperative surgical decision-making as well as to identify molecular biomarkers or therapeutic targets for the drug-resistant epilepsies.

## Introduction

The ability to map the transcriptional landscape of human epilepsy *in vivo* remains a major challenge in developing our understanding of the focal epilepsies. To date, the primary methods of profiling human epileptic brain tissue have been through fresh tissue samples removed during neurosurgery, and post-mortem studies (1). Stereo-electroencephalography (SEEG) involves implantation of intracranial electrodes into deep brain structures to characterise the epileptogenic network and guide surgical intervention in focal epilepsy (2,3). Typically, intracranial depth electrodes are explanted and disposed of upon completion of the SEEG evaluation. Recent studies have demonstrated the feasibility of retaining intracranial depth electrodes upon explantation and using the trace amounts of nucleic acids from fluid and cells adherent to the surface for identification of brain-specific somatic variants in malformations of cortical development (MCD) and neuronal migration disorders (4–10). Given that electrodes are implanted to provide coverage of a broad sample of deep and neocortical brain structures, this provides a unique opportunity to sample the human brain micro-environment in a minimally invasive manner. Furthermore, many of these electrodes are implanted in regions which are ultimately remote to the site of seizure onset, in tissue which is uninvolved in seizure onset or propagation (i.e. healthy brain). These samples represent a unique internal control which would normally be inaccessible *in vivo*, due to the technical and ethical barriers to sampling normal brain distant to epileptogenic tissue.

We recently reported the isolation of nucleic acids including RNA from SEEG contacts followed by genome-wide transcriptome and methylome profiling, overlaying this with clinical and neurophysiological data to achieve multimodal profiling of focal epilepsy (11). Profiles comprising up to one hundred million mapped reads per sample were obtained from individuals with focal cortical dysplasia (FCD), non-lesional temporal lobe epilepsy (TLE), and Rasmussen’s encephalitis (RE). There remain additional opportunities to understand the type and value of the sampled gene expression profiles on SEEG contacts. For example, to what extent do the recovered transcripts reflect the local cellular and electrophysiologic milieu? Are transcripts recovered from a broad range of cell types, including neurons, astrocytes and microglia? Is it possible to use such bulk RNA sequencing data to estimate cell type proportions and to do this throughout different brain regions? Indeed, do the data allow detection of brain structure-enriched genes including those in the hippocampus, middle temporal gyrus (MTG), cingulate, frontal lobe, along with grey and white matter? How well do gene expression profiles and the cell-type proportions correlate with epileptogenicity and the biological and pathological processes regulated by these genes? Finally, what are the current limitations with this approach? Here we present further analysis of the transcriptome data from this study. The findings suggest it is possible to integrate gene expression and neurophysiological data during the SEEG evaluation to better characterise the epileptogenic network.

## Methods

### Participant recruitment, sample collection and neurophysiological data acquisition

Patient information has been reported previously (11). Briefly, three patients with refractory focal epilepsy undergoing SEEG evaluation as part of the pre-surgical evaluation at the National Epilepsy Surgery Programme in Beaumont Hospital, Dublin, were recruited. Patient A had FCD type IIa, Patient B a non-lesional TLE and Patient C had RE. Between six and ten depth electrodes (Microdeep, DIXI Medical) were implanted under robot-assisted stereotactic guidance. Continuous EEG at 1024 Hz and concurrent video was acquired with the Xltek EEG System (Natus Inc). The duration of SEEG monitoring ranged from eleven to eighteen days. Upon explantation, electrodes were collected in sterile containers, immediately placed on dry ice and subsequently stored at -80°C. Groups of five adjacent electrode contacts, from the base or tip of the SEEG electrode, corresponding to anatomical regions as defined by post-implantation CT and MRI co-registration were clipped and pooled in each sample.

### RNA extraction, purification, and sequencing

Technical aspects of isolating and sequencing RNA from SEEG electrodes were previously reported (11). Briefly, RNA was isolated using the PicoPure RNA Isolation Kit (Thermo Fisher Scientific). RNA purification was performed using the GeneJET RNA Purification Kit (Thermo Fisher Scientific). The RNA integrity number (RIN) ranged from 1.2 to 5.9 for Patient A, 2.6 to 8.4 for Patient B, and 5.2 to 7.6 for Patient C. FLASH-seq, a single cell sequencing protocol suited for low input RNA sequencing was used for the bulk RNA samples (12). Raw FASTQ files were processed to remove adapter sequences using Cutadapt (v4.9) and were mapped against the human genome (GRCh38) using STAR aligner (13,14). The sample from the electrode placed within the FCD (SOZ) did not pass quality control, and so no seizure-onset samples were available from Patient A.

### Neurophysiological data analysis

Electroencephalogram recordings were analysed by an experienced Clinical Neurophysiologist. Each brain region sampled was designated as seizure-onset zone (SOZ), propagation zone (PZ), or non-involved zone (NIZ) by standard visual analysis. A channel was considered as PZ if it was involved within ten seconds of seizure onset. In addition, the Epileptogenicity Index (EI) plugin for Anywave (Marseille, France) was used to quantify the epileptogenicity of a given bipolar SEEG channel consisting of two adjacent grey matter contacts (15,16). Given that each sample consisted of five contacts, the channel with the maximum EI value in each anatomical region was matched with its corresponding pooled RNA sample.

### Transcriptome data analysis

Raw transcriptome counts were normalised, and differential gene expression analysis was performed using Deseq2 (Bioconductor)(17). ClusterProfiler (Bioconductor) was used for gene ontology (GO) enrichment analysis (18). The BRETIGEA R package was used for estimation of cell-type proportions from the bulk transcriptome data (19). For the identification of genes enriched in specific brain structures, we used the Allen Human Brain Atlas Microarray data explorer (20). The target region was contrasted with the region through which the SEEG electrode traverses during implantation en route to the target (e.g. the hippocampus was contrasted with the middle temporal gyrus). This list of transcripts was cross-referenced with the sample transcriptomes, and genes which were well represented in our data, significantly enriched, and with a high fold change were included.

### Statistics and data visualisation

Student’s unpaired t-test (two-tailed) and one-way ANOVA tests were performed using SciPy. Seaborn and Altair-Vega Python packages were used for data visualisation (21,22).

## Results

### Clinical characteristics

The clinical, pathological, and surgical outcome details were reported before (11), and are summarised in Supplementary Table 1 and Figure 1. Pathological diagnoses were through histological examinations by an experienced neuropathologist. Twenty-four SEEG electrode samples were selected to give broad representation of the SOZ, PZ, and NIZ. Study participants varied in terms of epilepsy aetiology, but also with respect to the extent of the epileptogenic network, from a focal frontal type II FCD (Patient A), to non-lesional TLE (Patient B), to right hemispheric RE (Patient C). Patient C was initially diagnosed with RE at sixteen years of age following frontal lobe biopsy and underwent early immunotherapy. At the time of SEEG evaluation, the immune component of the RE was thought to be quiescent, later confirmed by histopathology.

**Figure 1:**
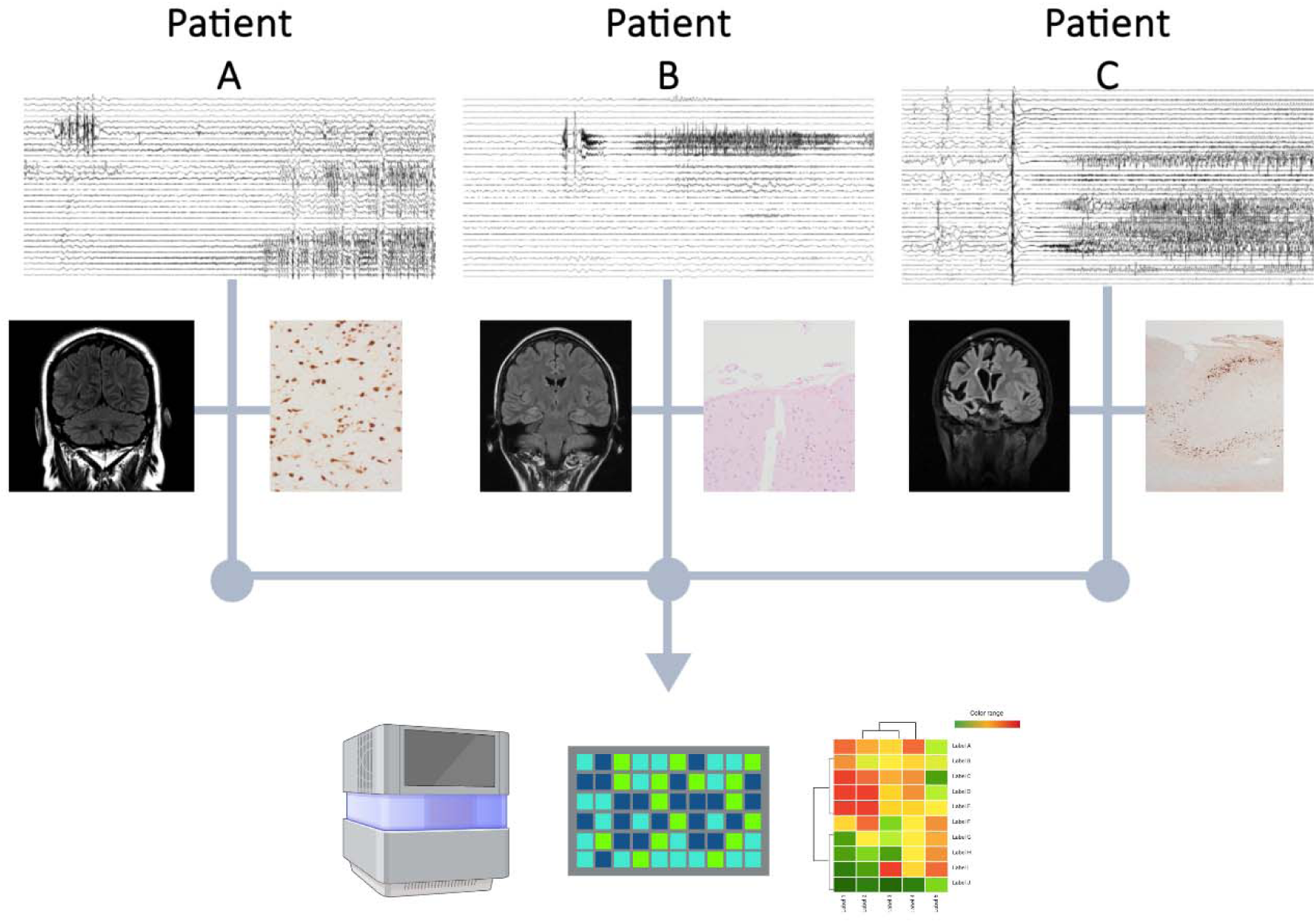
Clinical and pathological characteristics of study participants. Illustrative representation of multimodal data integration.

### RNA sequencing using surface material on explanted SEEG electrodes

RNA from 19 of 24 samples was successfully sequenced (11). Briefly, four SEEGs samples were achieved from Patient A with an average of 76 million transcriptome reads mapped to the human genome, while eight samples were achieved from Patient B and seven samples from Patient C (average ninety-seven million reads mapped). Principal component analysis (PCA) demonstrated that the greatest variance between transcriptome profiles was between patients rather than by anatomical origin or by involvement in the epileptogenesis (Figure 2A). Of note, two samples from Patient A were identified as outliers and were excluded from further analyses.

**Figure 2:**
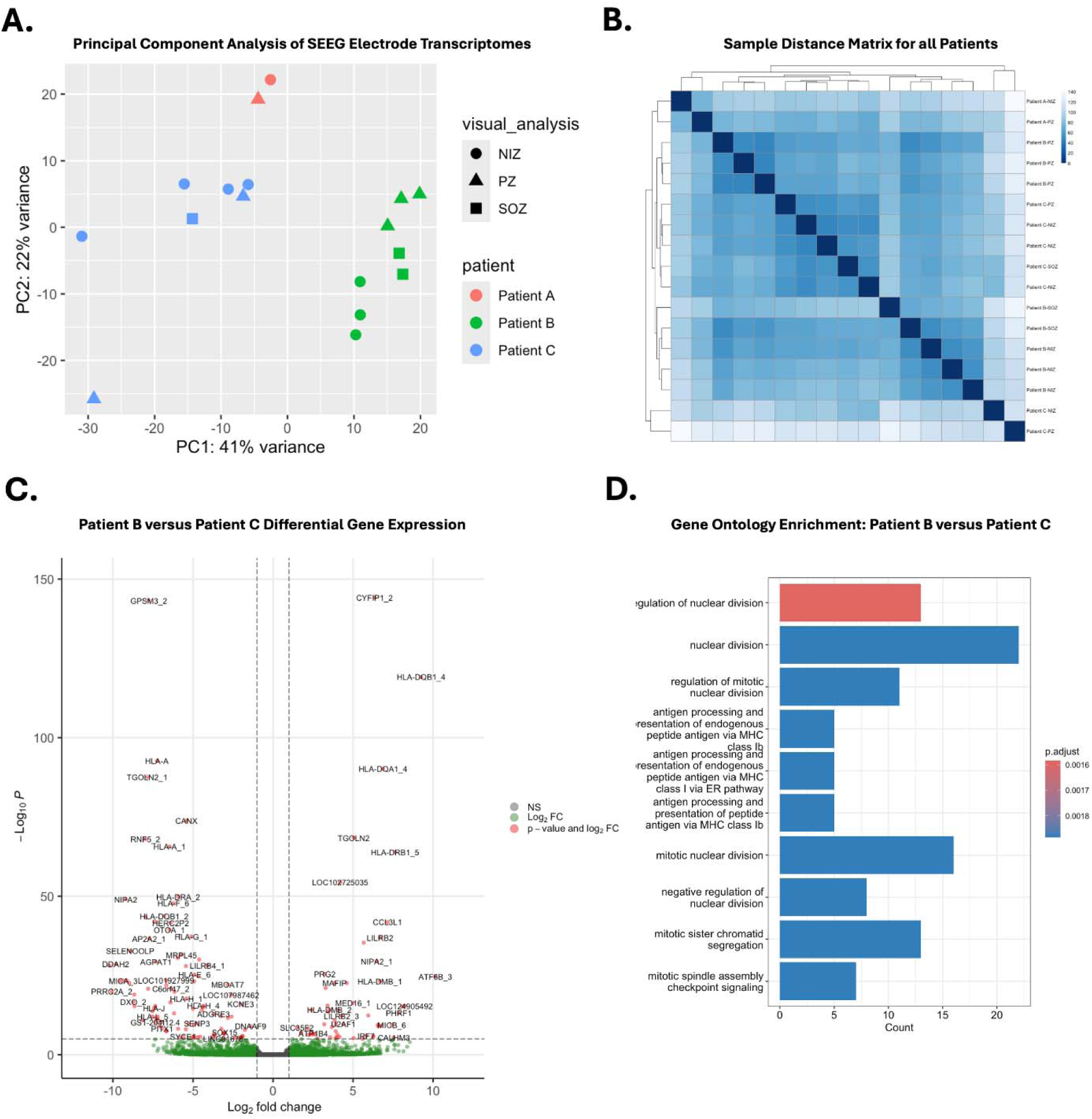
Transcriptome variation between patients. A. Principal component analysis showing variation between transcriptomes derived from each SEEG electrode sample. B. Sample distance matrix from all samples to all samples. C. Volcano plot of differentially expressed genes between Patient B and Patient C samples. D. Enriched biological processes in Patient B samples when compared to Patient C samples.

We first investigated the potential sources of the clustering of transcriptome differences between individuals. There was no significant difference in the expression of the commonly used housekeeping genes *GAPDH* and β*-Actin* across samples between patients (Supplementary Figure 1A). Among the most significantly differentially expressed genes between patients were human leucocyte antigen (HLA) related genes such as *HLA-DRB5*, *HLA-DRB1*, *HLA-DRB3-1*, and *HLA-DQA1* (Supplementary Figure 1B). These genes were expressed at consistent levels across samples from each patient. Patients B and C show high expression of *XIST* and *DDX3Y* consistent with female sex, while Patient A does not express these genes consistent with male sex. Gene ontology (GO) analysis found that the terms most enriched in Patient C versus Patient B relate to nuclear division and antigen processing and presentation (Supplementary Table 2).

### Transcripts from the major brain cell types are present on SEEG electrode surfaces

We previously reported the presence of both excitatory and inhibitory neuron marker genes among transcripts on SEEG contacts (11). To extend these insights, we sought to understand whether contacts carry transcripts from additional brain cell types. Accordingly, we explored the presence of different cell phenotype markers among the detected transcriptomes using the Allen Brain Atlas and ranked list of cell-type gene markers provided by McKenzie et al. selecting the genes with the highest specificity for a given cell type from human datasets(19). Cell-type specific gene markers for neurons, including excitatory and inhibitory neurons (*RELN*, *SNAP25*, *CNR1*, *SLC17A1*, *GAD1*, *LAMP5* and *VIP*), astrocytes (*GFAP, AQP4*), microglia (*CCL4, ITGAX*) endothelial cells (*APOLD1*, *VWF*), and oligodendrocyte progenitor cells (*PDGFRA*) were present in the transcriptomes derived from SEEG electrodes (Figure 3). Marker genes for each cell type were present across samples from each patient, and these cell types were represented irrespective of involvement of a sample in seizure onset or propagation. Some common marker genes such as *RELN*, *AQP4*, and *CCL4* were not detected across all samples. These findings demonstrate SEEG contacts retain gene expression data from a diverse cellular landscape from both normal and seizure-generating brain structures.

**Figure 3:**
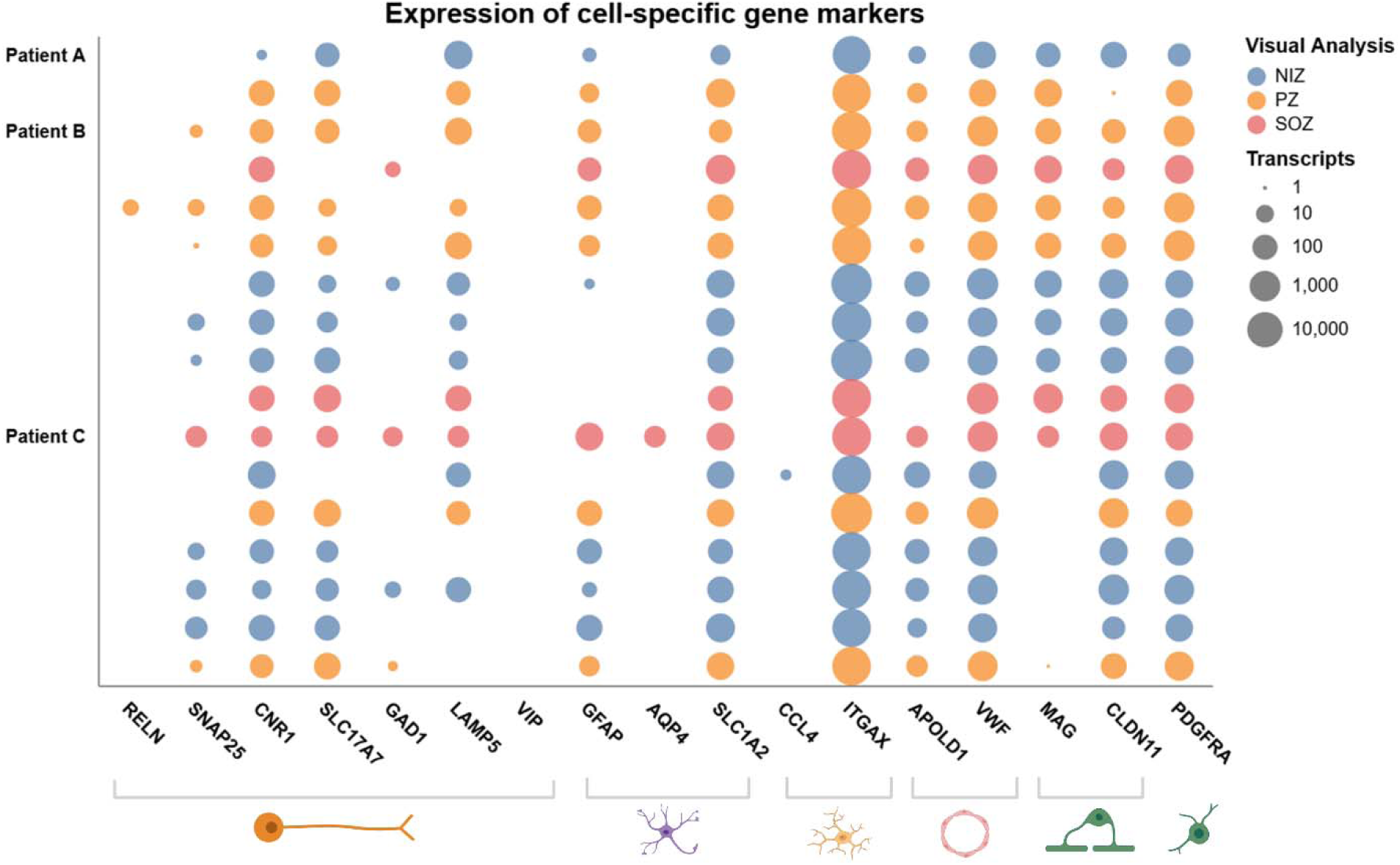
Range of cell-type specific transcripts identified from RNA sequencing. Dot matrix showing number of transcripts (logarithmic scale) of selected gene markers of neuronal and non-neuronal cell types present in each sample. Markers for neurons (*RELN*, *SNAP25*, *CNR1*, *SLC17A1*, *GAD1*, *LAMP5* and *VIP*), astrocytes (*GFAP, AQP4*), microglia (*CCL4, ITGAX*) endothelial cells (*APOLD1*, *VWF*), and oligodendrocyte progenitor cells (*PDGFRA*) are included. Samples are coloured according to the involvement in the ictogenic network.

### Cell type proportions estimated by bulk transcriptomes differ according to patient

To understand if electrode contacts reflect a proportional contribution of major cell types to the transcriptional signature, we next used the bulk transcriptome data to estimate the relative proportion of each major brain cell type (Figure 4). Due to the more limited extent of zone coverage from Patient A, we focused this analysis on Patients B and C only. This revealed representation of multiple major cell types for NIZ, PZ and SOZ as well as across patients (Figure 4A, B). The results indicated a similar ratio of neurons to other cell types (e.g. astrocytes). We noted a tendency to a higher proportion of oligodendrocytes in the SOZ when compared to the PZ and NIZ (p=0.1364) samples across all patients (Figure 4C) and found a significantly higher relative proportion of neuronal cells in the Patient B samples when compared to Patient C (p=0.0012) (Figure 4F). Last, we considered whether the placement of the electrode in terms of depth and brain region, had an influence on the cellular sources of recovered transcripts on contacts. However, cell type proportions were consistent across samples taken from deep and superficial structures (Figure 4D) and across anatomical regions (Figure 4E). Taken together, the findings confirm transcriptome profiles from SEEGs display specific contributions from cell types and may be informative about the local neuropathology at sites of implantation.

**Figure 4:**
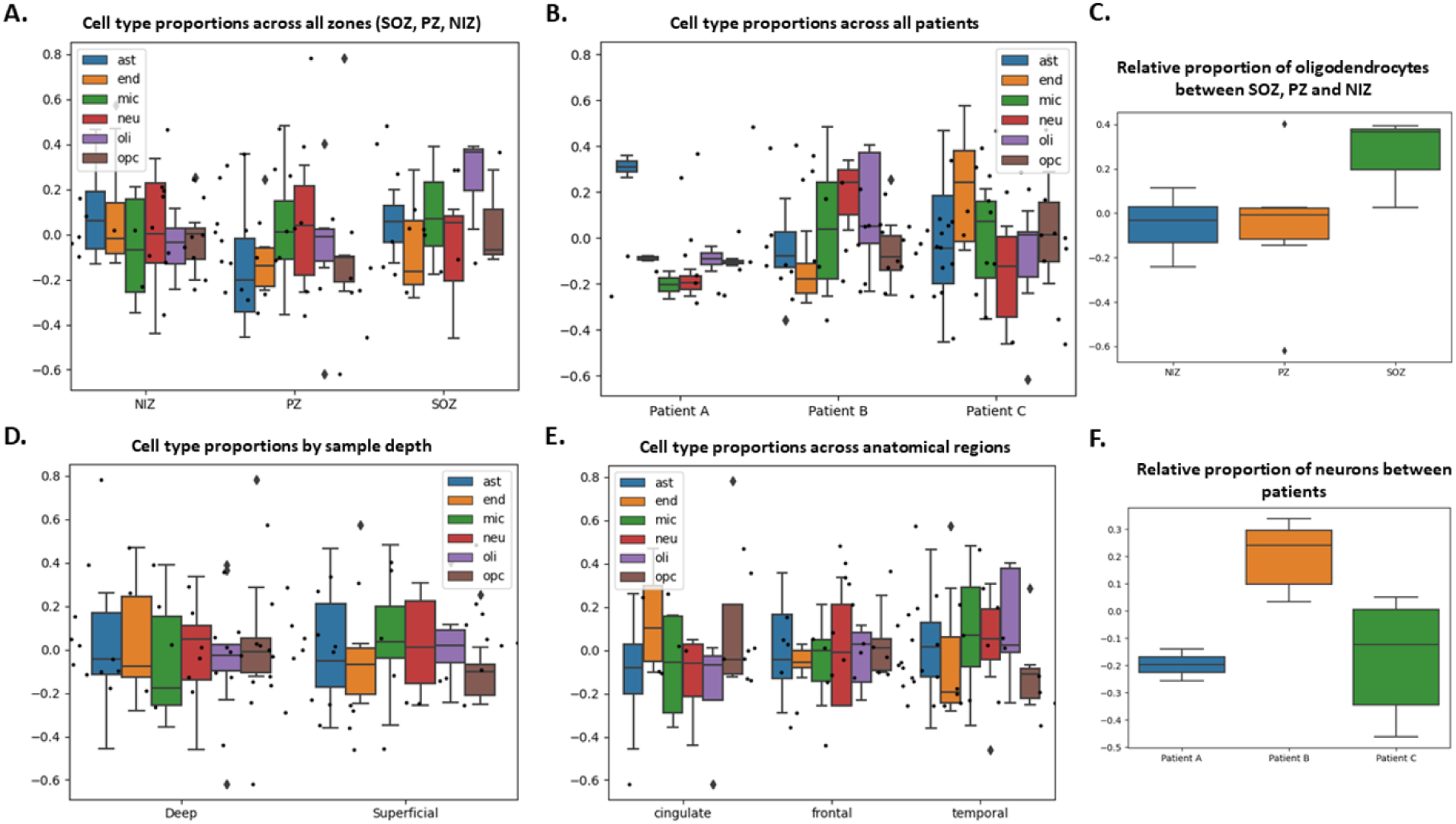
Cell type proportion analysis from bulk RNA sequencing. Relative proportions of brain cell types across samples using BRETIGEA. A. Cell type proportions between the NIZ, PZ, and SOZ. B. Cell type proportions across patients. C. Relative proportions of oligodendrocytes between the NIZ, PZ, and SOZ (p=0.1364, one-way ANOVA). D. Cell type proportions between samples implanted within deep subcortical structures and samples placed within superficial cortical structures. E. Cell type proportions in the frontal lobe, temporal lobe, and cingulate cortex. The single insula sample is excluded. F. Significantly higher proportion of neuronal cells between Patient B samples compared to Patient C (p=0.0012, one-way ANOVA).

### SEEG contacts display brain region molecular profiles from site of implantation

To complement the cell type analysis, we explored whether SEEGs retained transcripts reflective of the brain structure in which they were originally implanted. (Figure 5). We identified transcripts that are highly enriched in the human hippocampus (*LIPG*, *FZD7*, *ALPK1*, *ITGA4*, *NRP1*), frontal and temporal neocortex (*TESPA1*, *FRMPD28*, *RXFP1*, *CPNE9*, *ADTRP*), and cingulate (*GDA*, *NRGN*, *SLC30A3*). Genes enriched in grey matter (*PCLO*, *ATP282*, *MAP3K9*) and white matter (*RGS1*, *FA2H*, *SLC45A3*) were also identified. Expression of *ADTRP*, which is highly enriched in the frontal cortex compared to the cingulate cortex was significantly lower in the cingulate samples (p=0.00104). Expression of *SLC30A3*, which we found to be enriched in the cingulate, was more highly expressed in the cingulate (p=0.00444). Expression of *PCLO* and *ATP2B*, which are both enriched in grey matter structures compared to white matter structures, had significantly lower expression in the cingulate (p=0.0099 and p=0.01 respectively) (Supplementary Figure 2).

**Figure 5:**
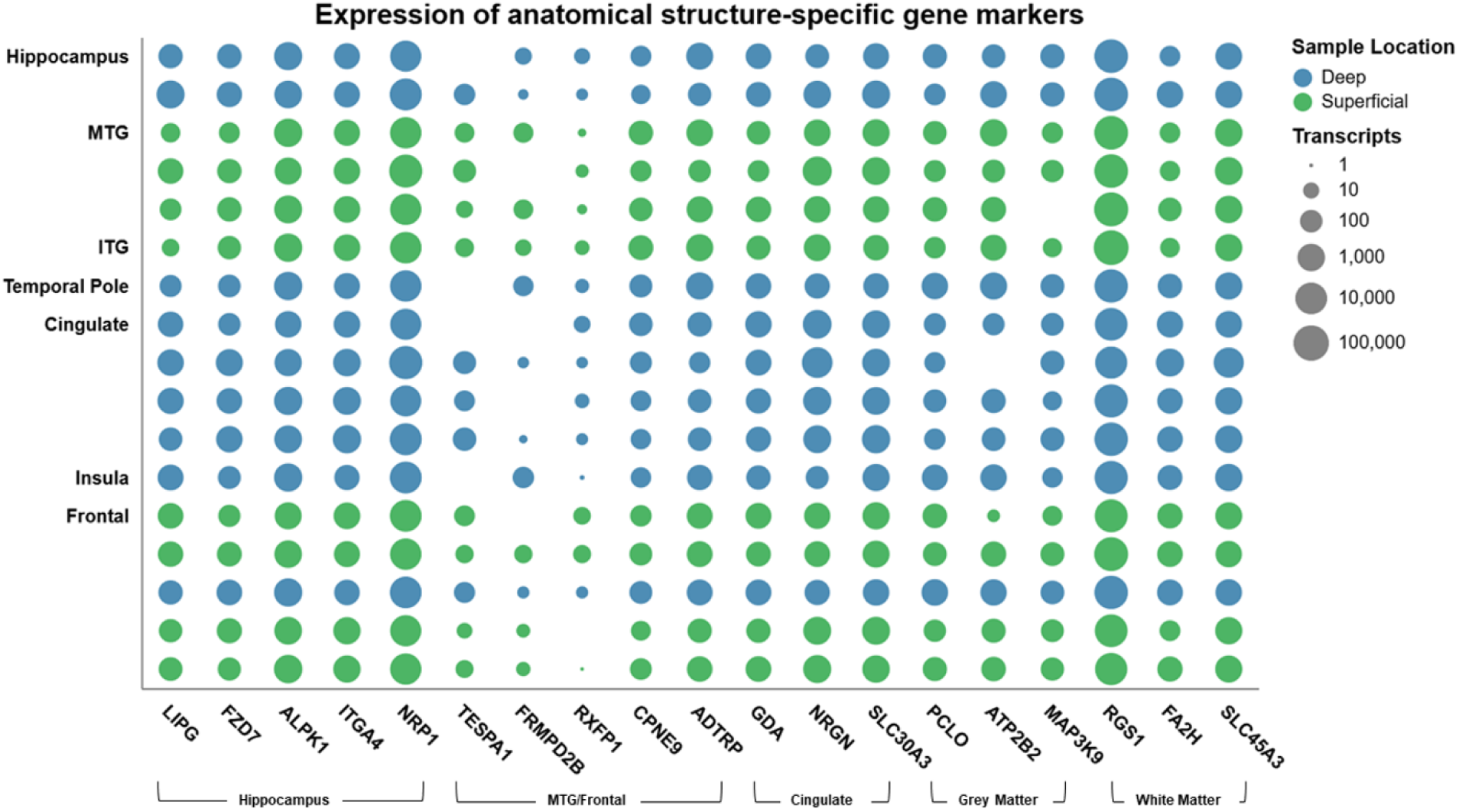
Expression of transcripts enriched in sampled brain structures across samples. Dot matrix showing number of transcripts (logarithmic scale) of selected gene found to be enriched in each brain region under investigation. *LIPG*, *FZD7*, *ALPK1*, *ITGA4*, *NRP1* represent genes enriched in the hippocampus. *TESPA1*, *FRMPD28*, *RXFP1*, *CPNE9*, *ADTRP* were enriched in the frontal and temporal neocortex, and *GDA*, *NRGN*, *SLC30A3* the cingulate cortex. Genes enriched in grey matter (*PCLO*, *ATP282*, *MAP3K9*) and white matter (*RGS1*, *FA2H*, *SLC45A3*) are also included. Blue hue represents samples from electrode contacts within deeper brain structures, and green those samples from contacts within the superficial cortical structures.

### Identification of gene activity reflective of epileptogenicity

We previously reported a number of differentially expressed genes for the three cases according to epileptogenicity index (11). To extend these insights, we investigated the presence of transcripts whose levels correlated most with locally recorded epileptiform activity (Figure 6A). Two protein-coding genes were differentially expressed across the seizure network zones: *TMEM130* and *PROSER3* (Figure 6B). While *TMEM130* was not detected in several samples, *PROSER3* was consistently lower in samples placed within SOZ and PZ when compared to the NIZ. This gene is poorly characterised with no known role in epilepsy. We did not detect differential expression of immediate early genes between SEEGs placed within high versus low epileptogenicity, as determined by visual analysis. GFAP expression was non-significantly increased in brain regions involved in seizure onset and spread (Supplementary Figure 1A). There was a non-significant decrease in *EGR1* across the SOZ, PZ, and NIZ (Supplementary Figure 1B).

**Figure 6:**
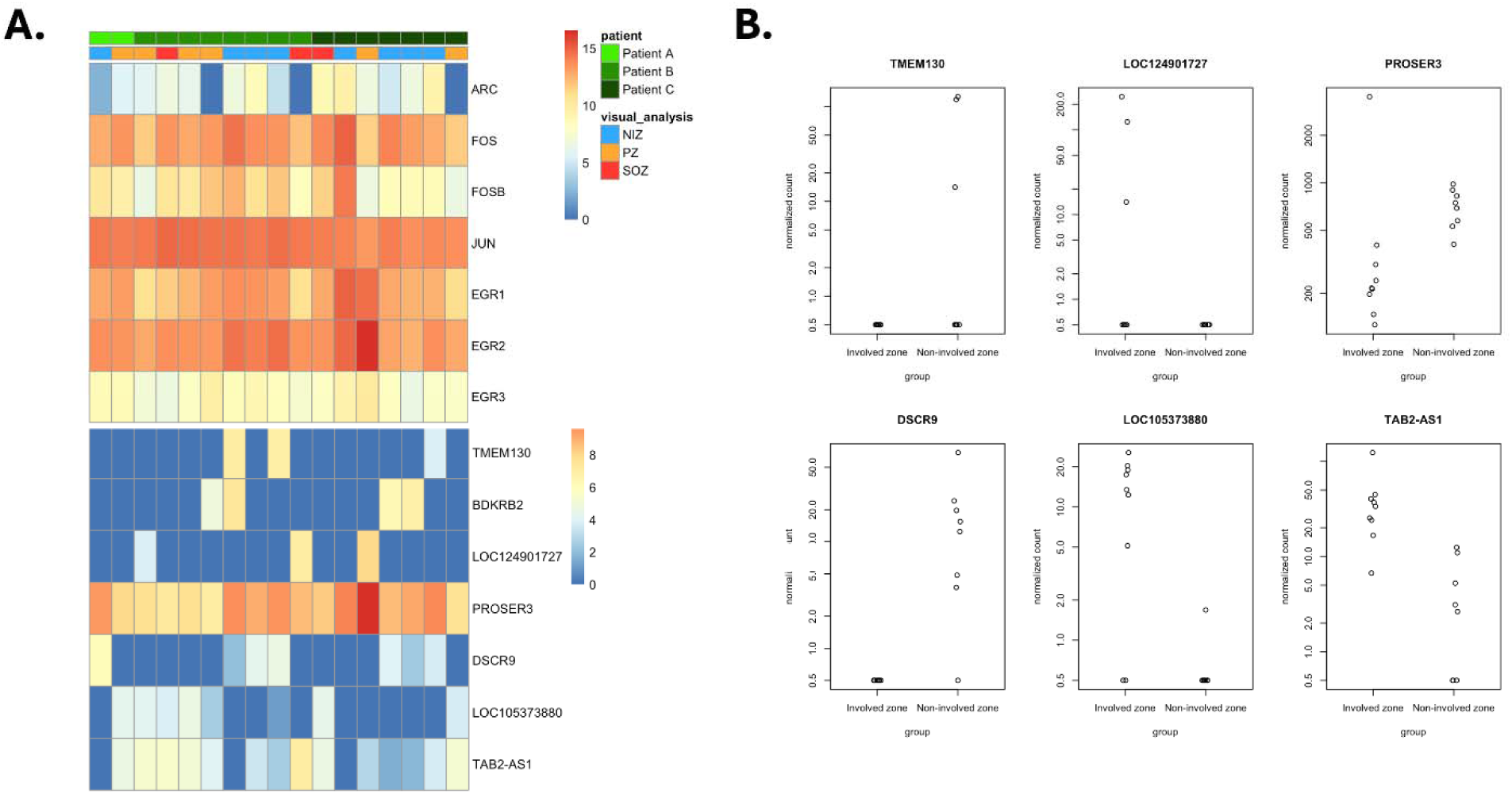
Correlation of gene expression with epileptogenicity. A. Heatmap of gene expression according to the patient and sample involvement within seizure network. The top panel displays genes expressed at higher levels and the bottom panel displays genes of lower expression levels. B. Scatter plots of expression of the significant differentially expressed genes between the involved zone (SOZ and PZ) and NIZ.

## Discussion

Focal epilepsy is a network disorder in which seizure activity generates and propagates from an epileptogenic brain region, such as a cortical dysplasia or heterotopia, across a wider group of brain structures (2,23). Genetic and computational EEG techniques have each advanced considerably in recent years and there now exists an opportunity to integrate both fields by combining neurophysiological data and molecular studies in order to elucidate the complex dynamics underlying seizure networks and ictogenesis. The present study extends the evidence that rich transcriptional data can be recovered from the surface of explanted depth electrodes from patients with epilepsy which reflects the local cellular and brain region environment. These data can provide insights into the molecular changes in zones of seizure onset and spread relative to healthy brain. Together, this approach offers new opportunities for understanding the molecular, cellular and network functions of the human brain in epilepsy.

Probing the human brain *in-vivo* remains a major challenge in neuroscience that has been partly unlocked by the recent demonstration that nucleic acids can be recovered from explanted depth electrodes (24). The first studies focused on DNA sequencing techniques using samples with lower spatial resolution. Montier et. al first reported using residual tissue on SEEG electrodes for genetic analysis, finding a somatic variant in *MEN1* from pooled electrode tips placed within a periventricular nodular heterotopia (4). Later, Ye et al. grouped whole electrodes, by broad anatomical location (quadrants), into three pools, across which a gradient of allele frequency in a *KCNT1* loss-of-function variant was noted to correspond to EEG findings (5). Checri et al. also identified mechanistic target of rapamycin (mTOR) pathway variants in patients with MCD but noted that spatial resolution was limited, as samples consisting of a single whole depth electrode which traversed neocortical, white matter, and deep cortical structures (6). These studies represent an important advance in our ability to characterise brain-specific somatic mutations in a minimally invasive manner, but we have yet to realise the potential impact on clinical practice and our understanding of the neurobiology of the epilepsy microenvironment. Indeed, measurement of DNA mutations or even epigenetic modifications alone cannot reveal whether these genes are actively being transcribed.

Here, we extend our recent demonstration that explanted SEEG contacts carry sufficient RNA to support high quality sequencing at high spatial resolution (11). The first finding in the present study on the same three cases with distinct aetiologies was that transcriptional signatures varied more between patient than by any other factor. This indicates SEEG profiles are highly individual-specific, likely reflecting the differences in underlying pathophysiology as well as genetic, developmental and temporal biochemical differences between individuals. Notably, although SEEG implantation plans are tailored to each individual patient, many of the samples included in this study were derived from a common set of brain regions in each patient. Indeed, samples from each patient showed a distinct pattern of expression of HLA-associated genes. This is expected, given that individuals display varied HLA types which are influenced by factors such as inheritance and ethnicity (25). The findings provide a technical validation of the ability to characterise neuro-immune markers throughout the brain that may provide insights to determine an individual’s susceptibility to disease and predict response to therapies.

Previous analysis of explanted electrodes did not explore the complete representation of cell types among the recovered nucleic acids (11). We found that we could use bulk RNA sequence data to estimate cell types and proportions throughout different brain regions *in vivo.* This revealed recovery of a mix of transcripts from excitatory and inhibitory neurons, consistent with our previous study (11), as well as contributions from the major classes of glia. Glial cells are important contributors to both epileptogenesis and ictogenesis (26–28). Thus, we extend the utility of electrode-recovered transcriptomic data and show value beyond simple differential gene expression analysis. Another important finding is that traces of the transcriptional signature of the brain region from which the electrode was removed are detectable. This counters the argument that the processes of removing the electrode from its implantation site would lose signal of origin during the passage through other regions on its route out of the brain. Electrodes remain in place for multiple days after implantation and cells may become adherent to the electrode surface. Precise information on the placement of each electrode contact within the brain allows direct correlation with the locally recorded neurophysiological activity after explantation. The relatively lower proportion of neuronal cell types in the samples from Patient C compared to Patient B is congruent with neuronal cell loss and gliosis seen in late-stage Rasmussen’s Encephalitis (29). The enrichment of genes with molecular functions related to antigen presentation and mitotic division in Patient C samples compared to Patient B could also correspond to underlying the immune processes and gliotic changes. Cell type proportions were consistent across deep and superficial samples, which further supports that sample integrity is maintained after the procedure of electrode removal. Taken together, the present analyses reveal a rich and diverse cellular contribution to the gene expression signatures on SEEGs that may provide insights into the pathophysiology of drug-resistant epilepsy and inform surgical decision-making.

This study demonstrates that the sample transcriptomes derived from the surface of SEEG electrodes contain a signal of the anatomical structure within which they were implanted. The expression of genes enriched in the cingulate and frontal cortex differed significantly depending on the sample position within the brain. We did not see a difference in expression of frontal/neocortical enriched genes between these structures and the cingulate. Furthermore, the expression of two grey matter enriched genes was lower in the cingulate samples. These findings may represent the passage of the electrode through anatomical structures during explantation, where deeper samples travel through the electrode tract, traversing white matter and superficial structures. While there is an inherent limitation in using gene markers to identify anatomical structures due to a lack of specificity, this was mitigated by selecting genes with the greatest change in expression between the two structures.

The ability to sample the nucleic acid and epigenetic landscape of SEEG contacts provide an opportunity to probe the mechanisms which sustain tissue that generates seizures (SOZ) while other connected sites to which seizure activity propagates (i.e. PZ) do not develop into tissue capable of initiating seizures. PZ contacts may bear molecular profiles in common with certain brain resilient states. It is well established experimentally that insults to the brain, when applied close to but below threshold for actual injury can activate endogenous protective programmes such that damage from a subsequent prolonged insult is reduced (‘tolerance’)(30). Notably, while repeated seizures can foster epileptogenic tissue (i.e. kindling), seizure activity that is subthreshold for injury can also activate neuroprotective programs in the brain that attenuate epileptogenesis (31). The tolerant brain is associated with changes to DNA methylation and down-regulation of pathways associated with calcium signaling and transport, energy and metabolism, among others (32,33). Notably, we detected lower expression of genes on SEEG contacts explanted from the PZ compared to SOZ for our TLE case associated with metal ions (e.g. multiple metallothionein genes), bioenergetics (*MTCH1/2*, *MTMR2/3/4*), as well as increases in processes associated with synaptic pruning (*C1QB*, *C1QC*) (11). Future studies may explore whether such patterns reflect protective adaptations, for example that limit spread of the SOZ, or are neutral or maladaptive to the nature of the seizure activity occurring within PZ tissue. The findings may inform our understanding of how networks may expand, resist epileptogenesis or resolve in individuals to inform diagnostic, biomarker and therapy directions.

There are limitations to consider in the present findings. The study was not powered or intended to draw firm conclusions regarding the biological mechanisms underlying focal epilepsy but rather support the feasibility of a method to gain insight to gene activity with cellular resolution integrated with neurophysiological data in the living human brain in a neurosurgical setting. The present study identified potential limitations with causality between gene activity detected on the surface of electrodes and neurophysiologic events. Specifically, we failed to detect higher levels of activity-regulated transcripts in SOZ and PZ compared to NIZ samples. Limited sample size and distinct aetiologies notwithstanding, immediate early genes such as *ARC*, *FOS*, and *JUN* do not appear to show higher expression in the SOZ than other sites. We must remember however, that these transcripts are typically upregulated within minutes of a seizure before rapidly declining. Surprisingly, *EGR1* showed a trend to graded reduction in expression across the NIZ, PZ, and SOZ. Immediate early genes undergo a rapid, transient increase in expression lasting for approximately thirty to sixty minutes after seizure activity(34). The process of reintroducing anti-seizure medications and anaesthetic induction in the operating theatre may also affect activity of these and other epilepsy-associated genes. However, the use of SEEG contacts allows for sampling of brain regions distant to the seizure focus, which may provide important insights into the dynamic changes throughout the brain that occur in response to seizures and epilepsy. Nevertheless, this study identifies several genes which showed a significant change in expression across the seizure network zones, indicating that the RNA sequencing data does contain a signal of epileptogenicity. Larger studies of people with epilepsy of similar aetiologies will be required to fully characterise the molecular signature of the seizure network. Future studies might also focus on whether a single contact carries sufficient material for RNA sequencing, thus providing a spatial resolution matching that of individual SEEG electrode channels. Additional research is also needed to determine the spatial extent of molecular findings across different pathologies and the effect of electrode explantation on sample integrity.

## Conclusion

In summary, the present study demonstrates that explanted SEEG contacts can be used to detect trace RNAs that reflect gene activity, cellular source and brain region of implantation. We also identified potential limitations in how well transcript information can be mapped to recorded seizure activity. Larger cohorts will be required to explore the extent to which this approach can inform our biological understanding of the pathophysiologic basis of epilepsy or inform surgical decision-making. Given the rapid advances in intraoperative molecular profiling of brain tumors to inform diagnosis and prognosis (35,36), we foresee applications of SEEG profiling integrated with neurophysiology for in-theatre diagnostic decision-making that could be applied to the epilepsies, as well as prospects for new biomarker and therapeutic target identification.

## Author contributions

JL conceived the study, performed data analysis and co-wrote the manuscript. AKD, AM and AS performed data analysis. VT and PWW supervised and co-conceived the study and edited the manuscript. DCH obtained funding, co-conceived the study and co-wrote the manuscript. All authors read and approved the final manuscript.

## Availability of data and materials

The RNA sequencing data are available from the gene expression omnibus (GSE268714). The rest of the data generated or analyzed during this study are included in this published article and its supplementary information files.

## Declarations

The present study was reviewed and approved by the Beaumont Hospital Medical Research Ethics Committee under study no. 20.58. All patients provided written informed consent.

## Acknowledgements

The authors thank the patients for the donation of samples for the present study. We thank Karina Halley for support with ethics and Austin Lacey for technical support with sample collection. We thank Kieron J. Sweeney and Donncha F. O’Brien for provision of neurosurgical samples and Ronan Kilbride, Department of Clinical Neurophysiology, Beaumont Hospital, Dublin for additional visual inspection of SEEG data. We thank Alan Beausang, Department of Neuropathology for the provision and interpretation of neuropathology slides and our other clinical colleagues at Beaumont Hospital. We thank Michael Dolan for advice on microglial genes. We thank the iCLOUD team at SDU and HPC team at Queen’s University Belfast (QUB) for computer and data management and the QUB Genomics facility.

