## Supplementary Figures for "Transcriptomic profiles from stereo-EEGs reveal the local cell microenvironment in human epilepsy"

Supporting Information

Transcriptomic profiles from stereo-electroencephalography electrodes reveal effects of epileptiform activity on the local cellular microenvironment in the human brain

Julian Larkin, Anuj Kumar Dwivedi, Arun Mahesh, Albert Sanfeliu, Vijay Tiwari, Peter Widdess-Walsh, and David C. Henshall

| Supplementary Table 1: Clinical characteristics of patients | | | | |  |  |
| --- | --- | --- | --- | --- | --- | --- |
|  | **Age** | **Sex** | **Epilepsy Classification** | **Epilepsy Duration (years)** | **No. ASMs** | **Surgical Outcome** |
| Patient A | 49 | Male | **Clinical:** Frontal lobe epilepsy  **Imaging:** Focal cortical dysplasia  **Pathology:** FCD IIa | 34 | 10 | 1A |
| Patient B | 40 | Female | **Clinical:** Temporal lobe epilepsy  **Imaging:** Non-lesional  **Pathology:** Chaslin’s subpial gliosis/non-specific | 8 | 7 | 2A |
| Patient C | 24 | Female | **Clinical:** Right hemisphere epilepsy  **Imaging:** Right hemi-atrophy, gliosis  **Pathology:** Gliosis, minimal T-cell infiltrate, perivascular chronic inflammation | 8 | 12 | 1A |

**Supplementary Table 2: Top 20 GO Enriched Processes, Patient B vs C (upregulated)**

| ID | Description | GeneRatio | BgRatio | RichFactor | FoldEnrichment | zScore | pvalue | p.adjust | qvalue | Count |
| --- | --- | --- | --- | --- | --- | --- | --- | --- | --- | --- |
| GO:0051783 | regulation of nuclear division | 13/284 | 151/18986 | 0.08609272 | 5.7554799 | 7.22970299 | 4.53988579318626e-07 | 0.00158397 | 0.00134524 | 13 |
| GO:0000280 | nuclear division | 22/284 | 451/18986 | 0.04878049 | 3.26107867 | 5.98863843 | 1.30614837769494e-06 | 0.0018843 | 0.00160031 | 22 |
| GO:0007088 | regulation of mitotic nuclear division | 11/284 | 121/18986 | 0.09090909 | 6.07746479 | 6.90449064 | 2.058294633187e-06 | 0.0018843 | 0.00160031 | 11 |
| GO:0002476 | antigen processing and presentation of endogenous peptide antigen via MHC class Ib | 5/284 | 17/18986 | 0.29411765 | 19.6623861 | 9.48614714 | 3.86109282371329e-06 | 0.0018843 | 0.00160031 | 5 |
| GO:0002484 | antigen processing and presentation of endogenous peptide antigen via MHC class I via ER pathway | 5/284 | 17/18986 | 0.29411765 | 19.6623861 | 9.48614714 | 3.86109282371329e-06 | 0.0018843 | 0.00160031 | 5 |
| GO:0002428 | antigen processing and presentation of peptide antigen via MHC class Ib | 5/284 | 18/18986 | 0.27777778 | 18.5700313 | 9.19006243 | 5.28089050143574e-06 | 0.0018843 | 0.00160031 | 5 |
| GO:0140014 | mitotic nuclear division | 16/284 | 282/18986 | 0.05673759 | 3.79302767 | 5.82308313 | 5.80177393094869e-06 | 0.0018843 | 0.00160031 | 16 |
| GO:0051784 | negative regulation of nuclear division | 8/284 | 67/18986 | 0.11940299 | 7.98234181 | 7.05520866 | 6.89811029668924e-06 | 0.0018843 | 0.00160031 | 8 |
| GO:0000070 | mitotic sister chromatid segregation | 13/284 | 194/18986 | 0.06701031 | 4.47978075 | 6.00325358 | 7.51547121139626e-06 | 0.0018843 | 0.00160031 | 13 |
| GO:0007094 | mitotic spindle assembly checkpoint signaling | 7/284 | 49/18986 | 0.14285714 | 9.55030181 | 7.38489525 | 7.81029290391766e-06 | 0.0018843 | 0.00160031 | 7 |
| GO:0071173 | spindle assembly checkpoint signaling | 7/284 | 49/18986 | 0.14285714 | 9.55030181 | 7.38489525 | 7.81029290391766e-06 | 0.0018843 | 0.00160031 | 7 |
| GO:0071174 | mitotic spindle checkpoint signaling | 7/284 | 49/18986 | 0.14285714 | 9.55030181 | 7.38489525 | 7.81029290391766e-06 | 0.0018843 | 0.00160031 | 7 |
| GO:0098813 | nuclear chromosome segregation | 17/284 | 324/18986 | 0.05246914 | 3.50767258 | 5.61028195 | 8.35925942936167e-06 | 0.0018843 | 0.00160031 | 17 |
| GO:0035115 | embryonic forelimb morphogenesis | 6/284 | 33/18986 | 0.18181818 | 12.1549296 | 7.90324174 | 8.37962046705594e-06 | 0.0018843 | 0.00160031 | 6 |
| GO:0031577 | spindle checkpoint signaling | 7/284 | 50/18986 | 0.14 | 9.35929578 | 7.29341652 | 8.96658022427903e-06 | 0.0018843 | 0.00160031 | 7 |
| GO:0033046 | negative regulation of sister chromatid segregation | 7/284 | 51/18986 | 0.1372549 | 9.17578017 | 7.20447077 | 1.02613221486692e-05 | 0.0018843 | 0.00160031 | 7 |
| GO:0033048 | negative regulation of mitotic sister chromatid segregation | 7/284 | 51/18986 | 0.1372549 | 9.17578017 | 7.20447077 | 1.02613221486692e-05 | 0.0018843 | 0.00160031 | 7 |
| GO:0045841 | negative regulation of mitotic metaphase/anaphase transition | 7/284 | 51/18986 | 0.1372549 | 9.17578017 | 7.20447077 | 1.02613221486692e-05 | 0.0018843 | 0.00160031 | 7 |
| GO:2000816 | negative regulation of mitotic sister chromatid separation | 7/284 | 51/18986 | 0.1372549 | 9.17578017 | 7.20447077 | 1.02613221486692e-05 | 0.0018843 | 0.00160031 | 7 |
| GO:0000819 | sister chromatid segregation | 14/284 | 235/18986 | 0.05957447 | 3.98267905 | 5.66955122 | 1.29241046075847e-05 | 0.00211209 | 0.00179376 | 14 |

**Supplementary Figure 1: Expression of housekeeping genes and HLA-related genes across patients**

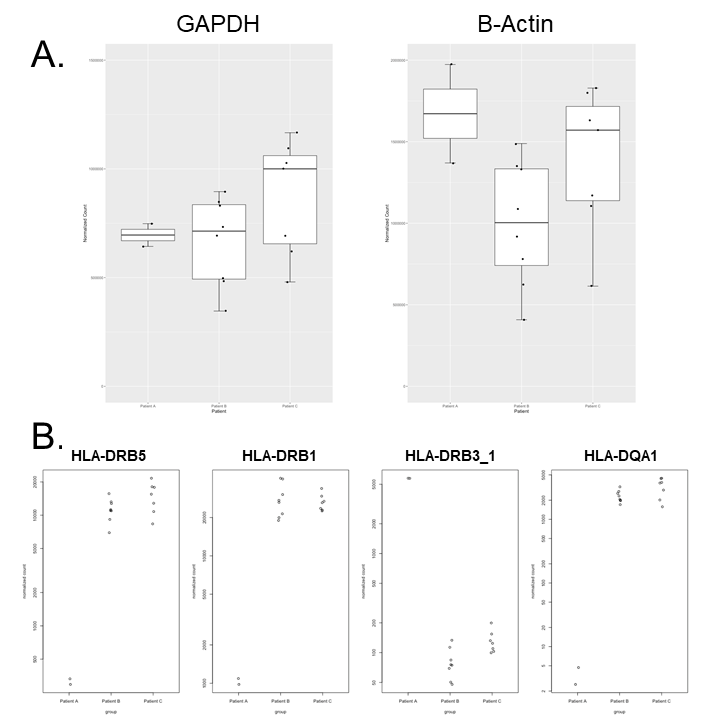

**Expression of housekeeping genes and HLA-related genes across patients:** A. *GADPH* and B-Actin expression does not significantly differ across samples between patients. B. *HLA-DRB5*, *HLA-DRB1*, *HLA-DRB3_1* and *HLA-DQA1* differ significantly between patients.

**Supplementary Figure 2: Expression of anatomical region-specific transcripts between the cingulate and all other structures**

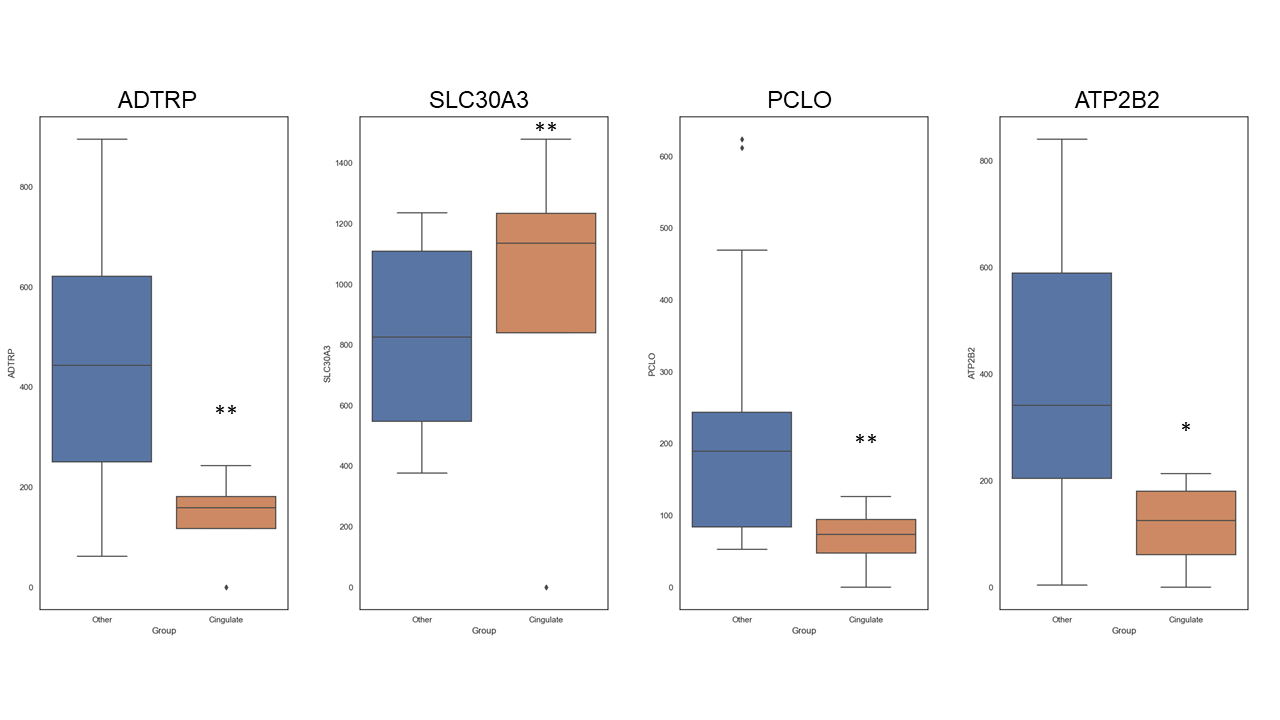

**Expression of anatomical region-specific transcripts between the cingulate and all other structures.** *ADTRP* (p=0.00104), *PCLO* (p=0.0099), and *ATP2B2* (p=0.01) expression is significantly lower in the cingulate than in all other structures. *SLC30A3* expression is higher in the cingulate compared to other structures (p=0.00444). Student’s unpaired t-test, two-tailed.

**Supplementary Figure 3: Normalised counts for selected genes across seizure network**

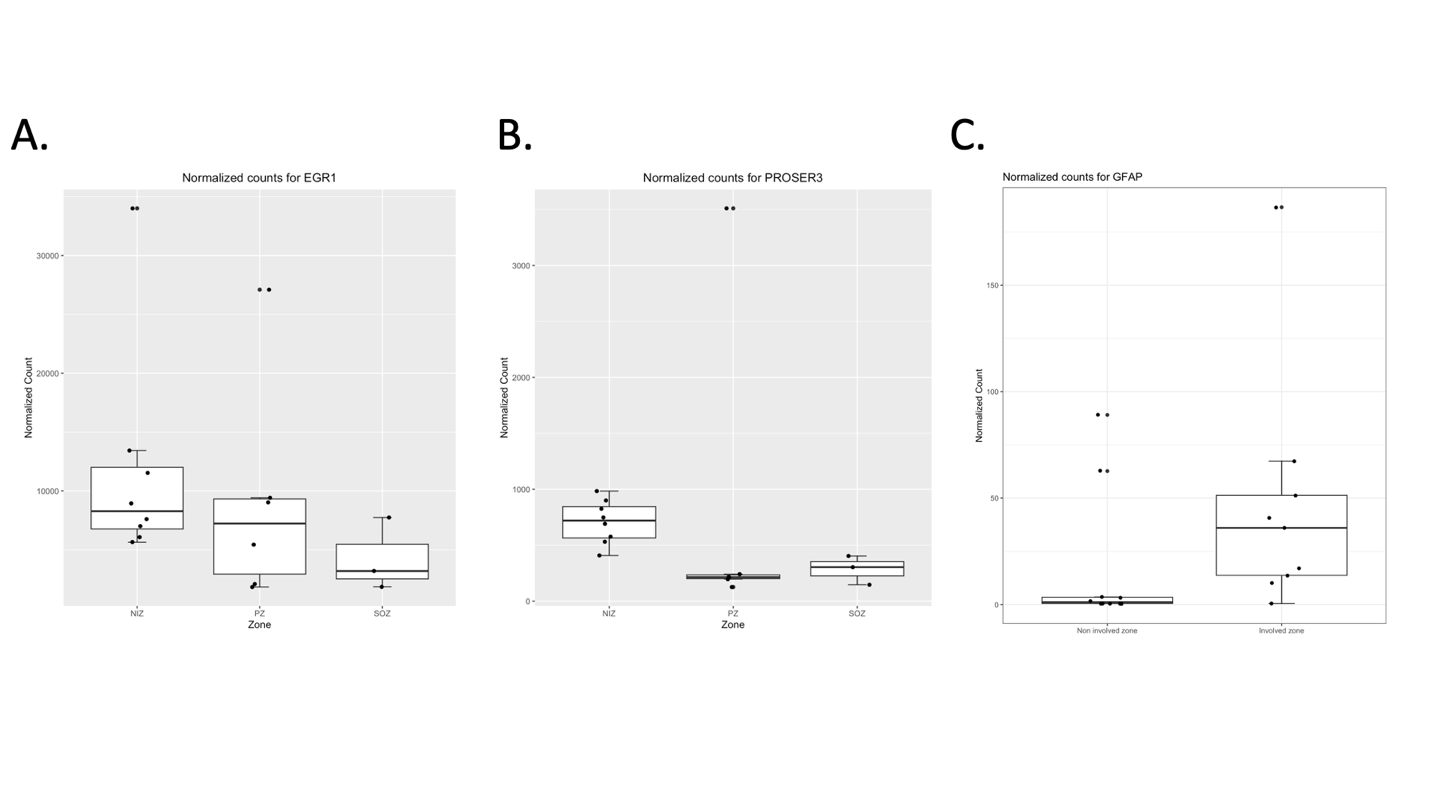

**Normalised counts for selected genes across seizure network:** A. Boxplot showing a non-significant decrease of EGR1 expression across the non-involved zone (NIZ), propagation zone (PZ), and seizure onset zone (SOZ). B. PROSER3 counts are significantly lower in the PZ and SOZ compared to the NIZ (p=0.0476). C. GFAP expression in the NIZ compared to the ictogenic network strucutres (PZ and SOZ).
